# A novel hybrid high speed mass spectrometer allows rapid translation from biomarker candidates to targeted clinical tests using 15N labeled proteins

**DOI:** 10.1101/2024.06.02.597029

**Authors:** Maria Wahle, Philip M. Remes, Vincent Albrecht, Johannes Mueller-Reif, Sophia Steigerwald, Tim Heymann, Lili Niu, Philip Lössl, Stevan Horning, Cristina C. Jacob, Matthias Mann

**Affiliations:** Department for Proteomics and Signal Transduction, Max-Planck Institute of Biochemistry, Martinsried, Germany; Thermo Fisher Scientific, San Jose, California; Proteomics Program, Novo Nordisk Foundation Center for Protein Research, Faculty of Health and Medical Sciences, University of Copenhagen, Copenhagen, Denmark; Absea Biotechnology GmbH, Berlin, Germany; Thermo Fisher Scientific (Bremen), GmbH, Bremen, Germany

## Abstract

Recent developments in affinity binder or mass spectrometry (MS)-based plasma proteomics are now producing panels of potential biomarker candidates for diagnosis or prognosis. However, clinical validation and implementation of these biomarkers remain limited by the reliance on dated triple quadrupole MS technology. Here, we evaluate a novel hybrid high-speed mass spectrometer, Stellar MS, which integrates the robustness of triple quadrupoles with the enhanced capabilities of an advanced linear ion trap analyzer. This instrument allows for extremely rapid and sensitive parallel reaction monitoring (PRM) and MS3 targeting. The Stellar MS allowed targeting thousands of peptides originally measured on Orbitrap Astral MS, achieving high reproducibility and low coefficients of variation (CV) as well as sensitivity and specificity sufficient for more than the top 1000 plasma proteins. Furthermore, we developed targeted assays for alcohol-related liver disease (ALD) biomarkers, showcasing the potential of Stellar MS in clinical applications. Absolute quantification is typically a requirement for clinical assays and we explore the use of ^15^ N labeled protein standards in a rapid, streamlined and generic manner. Our results indicate that Stellar MS can bridge the gap between proteomics discovery and routine clinical testing, enhancing the diagnostic and prognostic utility of protein biomarkers.

## Introduction

The field of mass spectrometry (MS)-based plasma proteomics has experienced remarkable advancements in recent years ^1–3^. Technological innovations and improvements in sample preparation workflows have expanded our ability to profile the plasma proteome in greater depth. This now enables discovery-based studies that consistently demonstrate the diagnostic potential of proteomics, which is starting to surpass the capabilities of current clinical assays.

Despite these advances, clinical laboratories predominantly continue to rely on antibody-based methods, such as ELISA, which focus on single protein targets. In contrast, targeted MS is routinely used in newborn screening, therapeutic drug monitoring, and hormone assays, which demonstrates that MS based methods can achieve regulatory and clinical acceptance.

Triple quadrupole mass spectrometers, known for their simple yet effective architecture, remain the gold standard for targeted protein biomarker measurements. These instruments provide high signal-to-noise ratios and exceptional robustness, essential for reliable clinical diagnostics and are readily used with isotope labeled standards for absolute quantification^4,5^. However, they are restricted to selected or multiple reaction monitoring (SRM/MRM), instead of comprehensive MS/MS spectra^6^. This limitation hampers the translation of proteomics discoveries into clinical applications, which typically takes many months even if sufficient specificity and sensitivity are achieved. Rare exceptions prove the point, as in the successful transfer of a targeted thyroglobulin quantification assay, which lacks reliable ELISA alternatives ^7,8^.

Liver disease is a growing public health concern given that more than one-third of the world’s population have steatotic liver disease, a substantial proportion of whom will progress to steatohepatitis, fibrosis and more severe, even life threatening conditions ^9–11^. As the disease is largely asymptomatic, cost effective and specific tests are urgently needed. In this context, we recently described protein biomarker panels with high prognostic and diagnostic potential for the diagnosis of different stages of alcohol-related liver disease (ALD), which matched or outperformed all of the test procedures in routine clinical use ^12^. However, these promising results have yet to be translated into standardized clinical tests.

Here we explore a novel hybrid high speed mass spectrometer (Stellar MS) to begin to address the above challenges. The Thermo Scientific Stellar MS is designed to combine the simplicity of triple quadrupoles with the advantages of an advanced dual-pressure linear ion trap as the final mass analyzer. In contrast to the last quadrupole of a triple quadruple, the linear ion trap captures the entire set of fragments of the peptide ions of interest, and then rapidly scans out their entire spectrum without loss. This results in a platform with familiar technologies, performing an experiment that is already well-accepted in the clinic, but at much increased scale and with the quantitative and qualitative benefits inherent to parallel fragment accumulation and full scan mass analysis. In the context of targeted mass spectrometry this promises unprecedented sensitivity, specificity and speed including parallel reaction monitoring (PRM) as well as MS3 capabilities.

Furthermore, we introduce the use of full-length ^15^N labeled protein standards in this context. Unlike peptide-based isotopic labeling, these standards control for variability in digestion and sample preparation, offering improved quantification accuracy. The ^15^N labeling also reduces potential interferences in MS2 spectra, increasing the reliability of fragment ion quantification^13^. This approach holds promise for detecting proteoforms and post-translational modifications, enhancing the diagnostic potential of protein biomarkers.

In this paper, we first contrast the Stellar MS to established triple quadrupole technology and then describe how to transfer peptide panels from the Thermo Scientific Orbitrap Astral instrument to targeted assays, making use of the Stellar MS’s adaptive real time retention time correlation. We find that essentially all Orbitrap Astral identified plasma peptides can successfully be targeted and describe how to prioritize the most promising ones. Finally, we employ a panel of ^15^N labeled protein biomarkers for liver disease to advance the prospect of a universal clinical liver disease test.

## Experimental Procedures

### Plasma sample preparation

Plasma samples were prepared based on the previously published plasma proteome-profiling pipeline^14^. Shortly, 45 µl of 100 mM Tris (pH 8.0) was added to 5 µl of plasma for a tenfold dilution and mixed thoroughly. 10 µl of diluted plasma were transferred into 10 µl of reduction/alkylation buffer (20 mM tris(2-carboxyethyl)phosphine (TCEP), 80 mM chloroacetic acid (CAA)) The digestion mix was shortly centrifuged up to 500xg and heated to 99 °C for 10 min, followed by cooling to room temperature for 2 min in a PCR cycler. For digestion of the denatured proteins, 20 µl of a freshly prepared digestion mix (0.025 µg/µl trypsin and LysC in ddH2O) was added for a final digestion volume of 40 µl, followed by incubation at 37 °C, shaking (1000 rpm) overnight. Following overnight digestion, the enzymatic activity was quenched by adding 60 µl of 0.2% trifluoroacetic acid (TFA). For mass spectrometry (MS) measurement, 500 ng (1 µl) or 250 ng (5 uL of a 1:10 dilution) of the digested peptides were loaded onto disposable Evotip C18 trap columns (Evosep Biosystems) according to the manufacturer’s instructions. Shortly, Evotips were soaked in 1-propanol and then activated with 0.1% formic acid (FA) in acetonitrile (ACN), centrifuging at 500xg for 2 min. This was followed by renewed soaking in 1-propanol and a washing step using 0.1% FA in water. To ensure proper sample loading, 100 µl of 0.2 % TFA was added to the Evotips and 1 µl of sample was added into the solution before centrifugation at 500xg for 3 min. Evotips were then washed with 0.1% FA before adding 150 µl 0.1 % FA to prevent drying of the C18 material. Evotips were kept at 4 °C till measurement and digested samples were stored at -20 °C.

### ^15^ N labeled protein preparation

^15^N labeled proteins were obtained from Absea Biotechnology GmbH (Berlin, Germany). The constructs were cloned into pET30a vectors and expressed as His-tagged proteins in E.coli BL21(DE3) cultured in ^15^N-supplemented M9 medium. The proteins were purified by immobilized-metal affinity chromatography and stored in phosphate-buffered saline (pH 7.4). Protein purity and absolute concentration were assessed by SDS-PAGE and amino acid analysis (AAA). For digestion, the proteins were diluted in a TEAB-based buffer (60 mM TEAB, 10 % ACN, 10 mM TCEP, 40 mM CAA). After denaturation, reduction and alkylation for 10 min at 74 °C and cooling down, trypsin and LysC were added at a 1:25 ratio and the proteins were digested overnight at 37 °C. The digestion was stopped by acidification to 1% TFA and the digest was either stored at -20 °C or directly loaded onto Evotips as described above. Digests were either loaded with 25 ng per digested protein, spiked in at 1 ng per digested protein or loaded in a 1:3 dilution series starting from 25 ng (described below). The precursor based labeling efficiency of each protein was estimated by extracting the MS1 raw intensities wherever a precursor was matched using alphaRaw. The extracted raw intensity ratio of the (M-1/M0) was compared to predicted raw intensity ratios for different ^15^N contents (from 100% to 95%) for each precursor generated using alphaBase.

### DDA and DIA LC-MS acquisition

Plasma sample single shots as well as the ^15^N labeled proteins were analyzed using the Evosep One liquid chromatography (LC) system (Evosep Biosystems) coupled to an Orbitrap Exploris 480 mass spectrometer or an Orbitrap Astral mass spectrometer (Thermo Fisher Scientific). Peptides were eluted from the Evotips with up to 35% ACN and separated on a 15 cm or 8 cm PepSep (Bruker Daltonics) column using the Evosep 30 samples per day (SPD) or 60 SPD method (44 min, 21 min) coupled to a 30 µm stainless steel emitter (Evosep). Data-dependent acquisition (DDA) data was only acquired on the Exploris 480 using a top15 method. The column temperature was maintained at 50 °C using a butterfly oven and the column was interfaced with an EASY-Spray Source (Thermo Fisher Scientific) with the emitter held at 2200 V. Full MS scans from 350-1400 m/z were acquired at a resolution of 60,000 at m/z of 200 with a normalized automatic gain control (AGC) target of 300% and a maximum injection time of 25 ms. MS/MS resolution was set to 15,000 (200 m/z) with a normalized AGC target of 200% and a maximum injection time of 22 ms. Only precursors with charge states between 2+ to 6+ were selected for sequencing and the precursor isolation windows were set to 1.3 Thomson (Th). Previously target precursors were excluded from (re-)sequencing for 30 s. The normalized collision energy for higher-energy collisional dissociation (HCD) fragmentation was set to 30%. Plasma single shots were acquired in data-independent acquisition (DIA) mode on both the Exploris 480 mass spectrometer as well as the Orbitrap Astral mass spectrometer as follows. For the Exploris 480 acquisitions, full MS scans from 350-1400 m/z were acquired at a resolution of 120,000 (200 m/z) with a normalized AGC target of 300% and a maximum injection time of 45 ms. For MS/MS scans, the collision energy was set to 30%, the resolution to 15,000 (200 m/z), the normalized AGC target to 3000% (Tune Version 3) and the maximum injection time to 25 ms. 49 equidistant DIA windows of 13.7 Th with 1 Th overlap were distributed across the m/z range of 361-1033. The Orbitrap Astral mass spectrometer was interfaced with an EASY-Spray source equipped with a FAIMS Pro device (all Thermo Scientific). A total carrier gas flow of 3.5 L/min was used and an electrospray voltage of 2.0 kV was applied for ionization with a FAIMS CV of -40 V. The MS1 spectra was recorded using the Orbitrap analyzer at 240k resolution (200 m/z) from m/z 380-980 using an automatic gain control (AGC) target of 500% and a maximum injection time of 3 ms. The Astral analyzer was used for MS/MS scans in data-independent mode with 3 Th non-overlapping isolation windows with a scan range of 150-2000 m/z. The precursor accumulation time was 7 ms and the AGC target was 500%. The isolated ions were fragmented using HCD with 25% normalized collision energy.

### Targeted LC-MS acquisition and method generation on Stellar MS

Targeted MS2 and MS3 assays were created for the liver disease assay by first analyzing neat solutions of the 12 proteins from Absea Biotechnology (see **Supplementary Table 1** below). Digested peptides from each protein were loaded onto 12 separate tips at 25 ng amounts and included 50 fmol of Pierce™ Peptide Retention Time Calibration mixture (PRTC, PN 88321). The peptides in each tip were analyzed with a DIA experiment on Stellar MS using 4 Th isolation windows, for both 60 and 100 SPD Evosep gradients. Within Skyline (skyline.ms) an indexed retention time (iRT) calculator was generated that used the PRTC retention times as experimental retention time anchors. In silico libraries for the peptide iRTs and spectra were created within the Skyline library generator tool that utilized a connection to the Prosit server to retrieve the libraries^15^. These libraries combined with the PRTC experimental retention times enabled facile selection in Skyline of the neat, heavy labeled peptides from each protein. Scheduled targeted assays with 1 min timed acquisition windows were then generated with PRM Conductor for 60 and 100 SPD gradients, and for MS2 and MS3 analysis, resulting in a total of four instrument methods. PRM Conductor uses a heuristic for selecting multiple MS2 fragments to become multiplex-isolated MS3 precursor ions (US patent 10665440B1). Briefly, the highest intensity product ions are selected that satisfy the criteria that they are either multiply charged, or if singly charged they should have mass-to-charge greater than the intact precursor. The scan rate for all methods was 125 kDa/s. The Q1 isolation windows were 1 Th and 2 Th for the MS2 and MS3 methods, respectively. The second isolation stage in the ion trap uses a calibrated isolation width that depends on the relative mass-to-charge values of the multiplexed MS3 precursors, and ranges from a minimum of 2 Th for the lowest m/z precursor in a set, up to a maximum of around 10 Th for precursors that are more than 4x the m/z of the lowest precursor ion. MS2 activation was HCD with normalized collision energy (NCE) 30%, while the second activation stage for MS3 was resonance CID with NCE 35%, 2 ms duration, and q = 0.22. The best combination of activation types for peptide MS3 analysis has yet to be resolved, and there are 4 combinations that can be used on Stellar MS, e.g. CID/CID, CID/HCD, HCD/CID, and HCD/HCD. Of note is that currently CID for the second activation stage is done serially for each product in a high-to-low-m/z fashion (US10665440B1) which has very high fragmentation efficiency and capture but costs an additional 2 ms per product ion.

Label-free targeted assays were created based on either Orbitrap Astral or Stellar MS peptide search results. The Orbitrap Astral DIA experiment was described above, while the Stellar MS’s DIA also used FAIMS with CV -40 V but used the gas-phase-fractionation (GPF) approach with isolation width 1 Th and 6 replicates covering the range 400-1000 Th, with precursor mass ranges m/z 400-500, 500-600, …, 900-1000. Peptide searching was performed with the CHIMERYS intelligent search algorithm (MSAID GmbH, Germany) in Thermo Scientific Proteome Discoverer Software 3.1 with fragment tolerance 0.5 Da. Search results for both Orbitrap Astral and Stellar MS were imported into Skyline, and PRM Conductor was used to create a set of targeted assays, where the “Keep all precs.” Option was selected, as we wanted to target every potential analyte, with the knowledge that many low concentration IDs could be unsuitable for a targeted assay. Adaptive RT alignment was utilized in the targeted assays based on reference data from the discovery runs. To create reference spectra for aligning to Orbitrap Astral discovery, the 3 Th isolation width spectra were combined in silico to create spectra having an effective 51 Th isolation width, using the “Combine DIA Windows for Reference” option in PRM Conductor. The Stellar MS GPF experiments already contained an extra set of 50 Th isolation width acquisitions to gather reference data. From the Orbitrap Astral IDs, first 8 targeted assays were created with two minutes scheduled acquisition windows. These 8 assays were run, and Skyline was used to find the most likely candidate peaks. A further four assays were then created for the same set of Orbitrap Astral ID’d peptides, but with 0.6 min acquisition windows to perform replicate injections. For the Stellar MS IDs, 3 assays were directly created from the discovery results, with 0.6 min acquisition windows. For both Orbitrap Astral and Stellar MS candidate targets, three replicates were acquired. The targets having CV greater than 30% were removed, and then PRM Conductor was used to select the precursors having at least 3 transitions satisfying the criteria of minimum absolute area 100, minimum signal-to-background ratio 2.0, minimum relative area 0.05, minimum time correlation to median peptide chromatogram 0.80, and LC peak width between 4.0 and 20.0 sec.

### Raw data analysis

The Orbitrap Exploris 480 raw files acquired in DDA mode were analyzed using Fragpipe 20.0 using the default workflow including or excluding ^15^N as a fixed modification for each amino acid. DIA raw files from both Orbitrap Exploris 480 and Orbitrap Astral were analyzed using DIA-NN 1.8 or CHIMERYS 2.0 without a spectral library. Raw files going directly into the targeted method generation were analyzed as described above. Targeted raw files were analyzed using Skyline-daily.

### Experimental design and statistical rationale

All experiments were done using human plasma obtained as described above. Altogether, the dataset including raw data files and search results was uploaded to MassIVE (see below). We used the same plasma batch for the benchmarking and technical evaluation. In brief, measurements with different gradients, input amounts were done in triplicates unless mentioned differently. The experimental design and statistical rational are described in the respective figure legends. Reproducibility and quantitative accuracy were evaluated using technical replicates. Proteins targeted in the ^15^N labeled assay were selected from a proposed biomarker panel described in a previous publication based on their production and purification efficiency.

### Development and analytical validation of targeted MS measurements

All experiments fall within the category of a Tier 3 targeted study. The data was acquired using PRM and transitions were selected based on experimental data. Quantitative information is retrieved using the summed peak areas as selected by Skyline.

## Results

### Principle of the novel hybrid mass spectrometer in comparison to a triple quadrupole MS (342)

The Stellar MS is a unique derivative of the Thermo Scientific Orbitrap Fusion Tribrid and TSQ Altis Plus mass spectrometer. It utilizes the Orbitrap Fusion Tribrid instrument control software, and the chassis, vacuum chamber, turbo pump, and quadrupole mass filter of the TSQ Altis Plus MS. The ion funnel, Q00 with low-pass-filtering capabilities, and dual-cell linear ion trap are identical to those used in the Thermo Scientific Orbitrap Ascend Tribrid MS, while the Q0 and Q2 multipoles were redesigned for Stellar MS. The linear ion trap rf system on the Stellar MS achieves a slightly higher frequency of 1260 kHz, up from around 1180 kHz. The dual-high-energy-dynode-single-multiplier, capable of single ion detection, used in the Orbitrap Ascend Tribrid MS is combined with a new photo-multiplier-based final detector stage that considerably increases the detector stability and longevity. This final detector stage is also used in the Orbitrap Astral MS.

Clinical targeted assays are typically designed for and run on triple quadrupole mass spectrometers. Conceptually, their layout is simple, starting with the precursor selection in the first quadrupole followed by fragmentation in the second quadrupole and product ion (transition) selection in the third quadrupole (**Figure1 A**). Given that only one fragment ion can be measured at a time, the number of transitions that can be measured at any given time is limited. Acquisition of full MS/MS spectra would be highly inefficient. Consequently, in MS assays only the most intense and more specific fragment ion are monitored, typically three to five per peptide.

**Figure 1:**
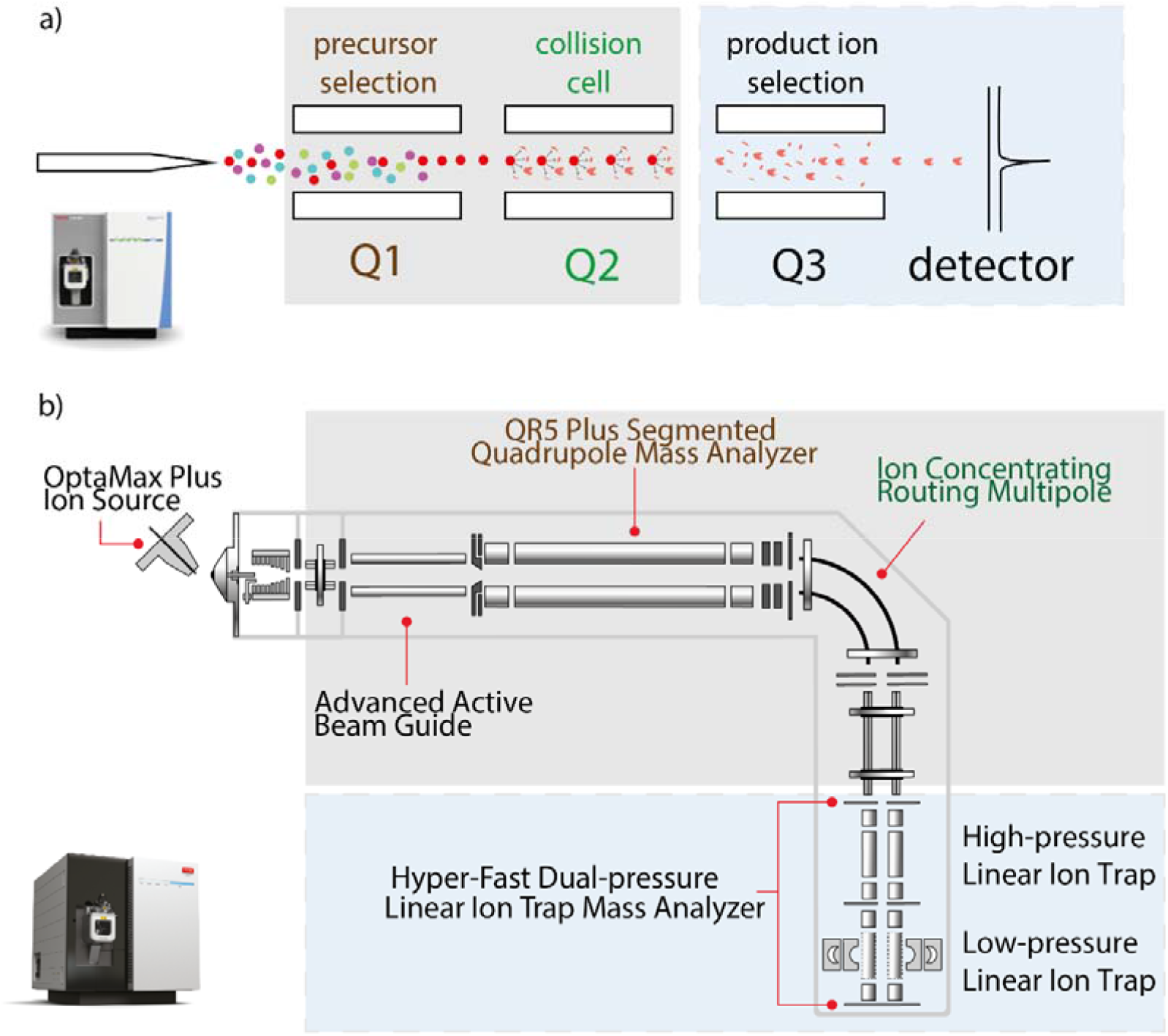
Comparison of traditional triple quadrupole mass spectrometer (MS) and Stellar MS. A Schematic of a traditional triple quadrupole MS, showing Q1 for precursor ion selection, Q2 for ion fragmentation, and Q3 for product ion selection. This setup limits the number of transitions to one (SRM) or a few, typically three to five (MRM). A photo of the Thermo TSQ Altis Plus is shown as an example of a triple quadruple instrument. B Schematic of the Stellar MS, featuring a dual-cell linear ion trap (LIT) instead of Q3. The LIT allows simultaneous ion accumulation and fragmentation, achieving acquisition rates of about 70 Hz and enabling MS3 analysis at up to 40 Hz. This enhances sensitivity and specificity compared to traditional SRM. Note that the only conceptual difference consists of replacing Q3 with the LIT (shaded blue in the figure), thus robustness and usability should be similar. The photo depicts the Thermo Stellar MS.

The novel hybrid high speed mass spectrometer Stellar MS follows similar design principles but instead of a third quadrupole it uses a dual-cell linear ion trap (LIT) as a mass analyzer **(Figure1 B**). Conceptually, the hardware and instrument control architecture are very similar to the Thermo Tribrid Series ^16^ but without the Orbitrap analyzer. Simultaneous ion accumulation/fragmentation in the collision cell with mass analysis in the LIT enables acquisition rates of about 70 Hz for typical peptide scan ranges of 200-1400 Th and allows ion injection to take place for ∼85% of the total acquisition time. The typical mass analysis scan rate is 125 kDa/s, which gives a peak full width half maximum of about 0.7 Th at m/z 622. The high pressure cell of the LIT is capable of performing waveform-based synchronous precursor selection multiplexing and resonance ion activation, which are particularly useful for MS3 analysis, which can be performed at up to 40 Hz. The dual-high-energy-dynode-single-multiplier with a photo-multiplier based final stage yields very high detector stability and longevity.

There are several advantages of parallel reaction monitoring (PRM) analysis performed on Stellar MS over the selected reaction monitoring (SRM) performed on a triple quadrupole. The first advantage is that multiple product ions of a precursor can be accumulated and then analyzed at once, instead of serially as in a triple quadrupole. For analysis of N product ions of a precursor that are formed at the same collision energy, the PRM analysis is at least N-fold more sensitive. For large scale assays, as SRM dwell times approach the approximate 1 ms needed for hardware switching between product ions, the advantage is greater than N-fold. The second advantage is that PRM analysis enables retrospective optimization of the transitions used for analysis, whereas SRM requires the transitions to be selected before the experiment starts. For molecules like peptides that typically fragment into many product ions, PRM effectively increases the selectivity of the analysis by allowing for post-acquisition transition selection.

We utilized a new software tool called PRM Conductor that takes advantage of the Stellar MS capabilities to more seamlessly build PRM assays (https://panoramaweb.org/prm_conductor.url). PRM Conductor is a Skyline external tool that consumes either DIA discovery data or PRM data. Its principal functions are to filter transitions with a set of simple criteria, such as absolute area, relative area, signal-to-background ratio, covariance to the median, and LC peak width. Precursors with at least a minimum number of qualifying transitions are accepted, typically three. Qualifying precursors are placed into the PRM assays and a user interface allows to visualize the effect of acquisition settings. Finally, the selected precursors can be exported to an instrument method file.

These features open up new possibilities for targeted assay design (**Figure 2 A**). Note that the Stellar MS instrument, can also acquire data in data independent acquisition (DIA) mode. From these nominal mass resolution datasets, preferably acquired by ‘gas phase fractionation (GPF) to increase depth’ ^17^, peptides deemed to be well targetable can be selected. Obtaining the targets on the same machine platform should reduce optimization iterations and thereby the time to design a targeted assay as the peptide is by definition well measurable. Alternatively, target lists could be transferred from discovery studies performed on different instruments. Here we were particularly interested in transfer from the recently introduced Orbitrap Astral MS. To this end, the peptides of interest are first evaluated in a set of targeted screens with up to 100 peptides per minute (see below and **Experimental Procedures**).

**Figure 2:**
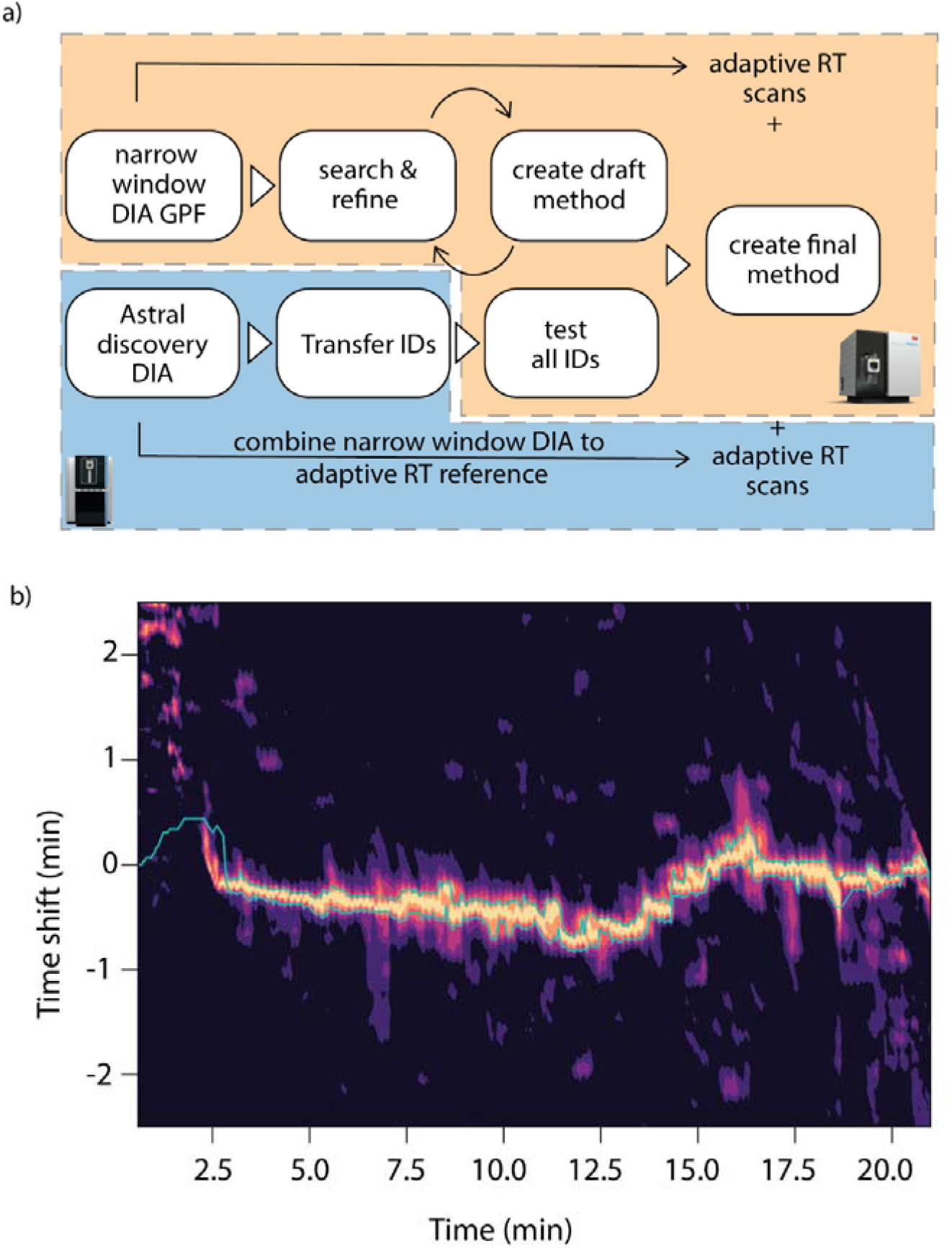
Development of targeted assays on Stellar MS using Orbitrap Astral discovery data. A Flowchart of targeted assay development for Stellar MS. Two pathways are shown: DIA on Stellar MS with gas phase fractionation (GPF) and transfer of Orbitrap Astral MS discovery data to Stellar MS. PRM Conductor software filters transitions, aiding high-quality precursor selection. B Contour plot of Adaptive RT cross correlations between alignment acquisition in a Stellar MS targeted run compared to the Orbitrap Astral MS in two different geographical locations. Retention time shifts within +/-30 seconds demonstrate robust transferability and accurate real-time alignment, enabling seamless transition from discovery to targeted assays across different platforms. The band indicated in green is the estimated retention time shift used as the basis for the adjusted acquisitions in the Adaptive RT scheme.

Subsequently, we set up targeted assays in a single step using PRM Conductor. This software interfaces with a new PRM data acquisition strategy called Adaptive RT that was described previously ^18^. Briefly, reference spectra from periodic acquisitions, such as from MS1 or DIA, are compressed and embedded in the instrument method file. These spectra can originate from discovery experiments on a Stellar MS, or another instrument such as an Orbitrap Exploris, or Orbitrap Astral. The PRM experiment includes these extra MS1 or DIA acquisitions and periodically compares their spectra with the reference data to update the scheduled acquisition windows in real time. This capability allows to reduce the scheduled acquisition windows to around a 1 min or less without suffering from chromatographic drift, and allows to more easily transfer discovery results to a PRM assay, including from one instrument to another.

Of note, the original discovery data was neither generated on the same human plasma samples nor on the same Evosep HPLC instruments but instead measured in Munich and San Jose. Nonetheless, retention time shifts of only +/-30 sec were observed which were readily recovered.

### Discovery DIA on Orbitrap Astral and translation of targets to Stellar MS

We compared the results of a single shot Orbitrap Astral discovery DIA measurement with a Stellar MS DIA gas phase fractionation for the selection of targetable peptides. The Orbitrap Astral measurement identified a broad range of peptides and protein groups. The peptides and proteins identified by the Stellar MS in DIA mode were largely a subset of these (**Figure 3A**). Note that the Stellar MS data were acquired with injection times twice as long as those used in the effectively fixed injection time Orbitrap Astral data (**Figure 3B**).

**Figure 3:**
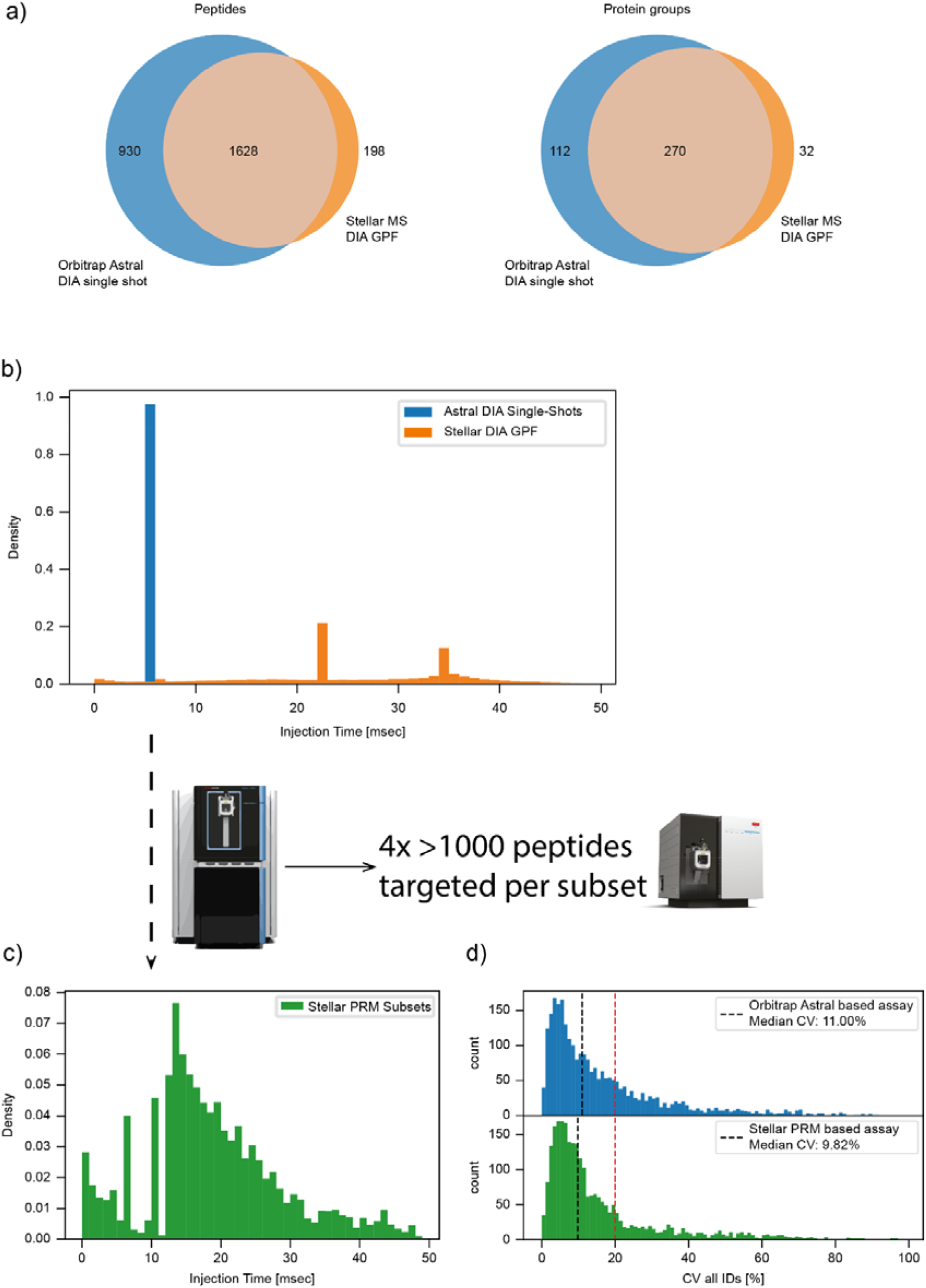
Discovery DIA on Orbitrap Astral and Stellar MS. A Peptides and protein groups identified in a single-shot Orbitrap Astral discovery DIA measurement and a Stellar MS DIA with gas phase fractionation (GPF). B Comparison of scan wise injection times for Orbitrap Astral single-shot DIA acquisition and Stellar MS DIA GPF. Stellar MS uses twice the injection time due to its restricted m/z range in each GPF run. C Scan wise injection times for Stellar MS parallel reaction monitoring (PRM) subset measurements of Orbitrap Astral discovery DIA data. Increased injection times and targeted measurements improve quantitative accuracy. D Coefficients of variation (CVs) for peptides measured in Orbitrap Astral DIA acquisition and Stellar MS PRM subset measurements. The targeted subsets on Stellar MS show a comparable median CV from 11 % to 9.82%, demonstrating the platform’s high reproducibility and sensitivity.

Next, we split the 2,558 Orbitrap Astral identified peptides into four subsets and remeasured those on the Stellar MS in a targeted manner. By increasing the injection times and spacing out the potential targets, we observed a significant reduction in the coefficients of variation (CVs). The targeted subsets achieved a comparable median CV of 11 % to 9.8 % (**Figure 3D**), underscoring the high sensitivity and reproducibility of Stellar MS and cross-validating peptide identifications between platforms.

We evaluated both targeted assay design modalities by designing a targeted assay for our neat plasma proteome measurements. This resulted in at least 40% more targetable peptides and at least 25% more targetable protein groups based on Orbitrap Astral data than Stellar MS data alone.

### Characterization of target lists from Orbitrap Astral and Stellar MS

To transition from a list of targetable peptides to a fully developed targeted assay, it is essential to select peptides that can be measured reproducibly in terms of retention time and signal intensity. As we wanted to move from selecting just a few peptides towards targeting thousands of peptides in a single assay, we needed to prioritize candidates from the discovery data in order to rapidly design our target list. In our first approach, we selected a minimum number of transitions per peptide, more specifically, we retained every peptide with at least three good transitions for computational testing. This filter was fairly efficient at eliminating peptides from Stellar MS DIA GPF data that lead to large CVs. However, it entailed a more than twice as high false negative rate on the Orbitrap Astral data (ratio of peptides eliminated although they were well targetable (**Figure 4 A, C**). We ascribe this to the fact that injection times in the Orbitrap Astral were comparatively short, resulting in lower quality transitions than the Stellar MS transitions with higher injection time. Likewise, this simple approach ignores edge effects in the gradient, namely the switch to a lower flowrate at 2.5 min and to high organic buffer at 20 min at the end of the gradient, making peptide retention times less predictable at those times.

**Figure 4:**
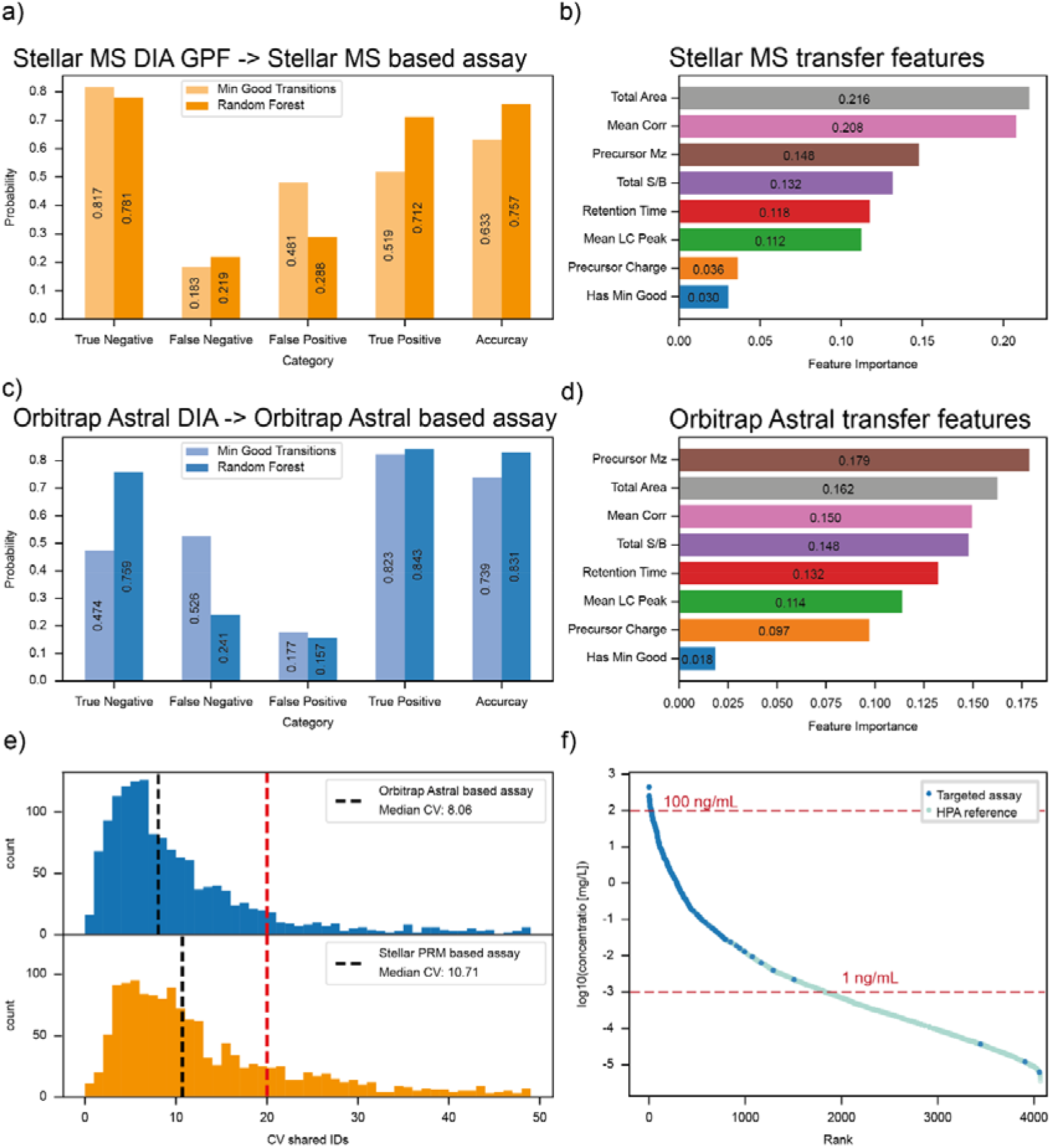
Comparison of Orbitrap Astral discovery DIA IDs and Stellar MS GPF IDs. A Performance metrics of a Stellar MS discovery-based PRM assay designed using a minimum number of good transitions (dark orange) compared to a random forest machine learning (ML) model (light orange). B Ranked feature importance for the Stellar MS-based random forest model used to design the PRM assay, highlighting the importance of total area, signal-to-noise ratio (S/B), and retention time. C Performance metrics of an Orbitrap Astral discovery-based PRM assay designed using a minimum number of good transitions (dark blue) compared to a random forest ML model (orange). D Ranked feature importance for the Orbitrap Astral-based random forest model used for the PRM assay design. E Coefficients of variation (CVs) for peptides measured in Stellar MS (orange) and Orbitrap Astral (blue) based PRM assays. Despite targeting more peptides in the Orbitrap Astral-based assay, the median CVs for shared peptides remained similar. F CV coverage of proteins reported in MS-based assays by the Human Protein Atlas (light blue) captured by the Stellar MS targeted assay based on Orbitrap Astral discovery data (dark blue), illustrating the broad dynamic range of protein abundances targeted.

We found that switching to a random forest classification of targetable peptides from discovery data improved the accuracy both starting from Orbitrap Astral data as well as from Stellar MS discovery data. Features like the total area, signal to noise ratio and mean area correlation turned out to have a higher feature importance than the minimum number of good transitions (**Figure 4 B, D**). Including more information and a machine learning approach thus promises of further improving the predictability of which peptides are well targetable from all those identified in discovery experiment setups.

We generated two targeted assays for Stellar MS, one based on Stellar MS discovery data (1826 peptides, 302 proteins) and the other on Orbitrap Astral DIA data (2558 peptides, 382 proteins) for a 21 min gradient. This means that we were able to target 40% more peptides and 25% more protein groups if we include the Orbitrap Astral discovery DIA data in the assay generation process.

When we compare peptides that are shared between both assays, the median CV of both assays was similar despite targeting many more peptides in the Orbitrap Astral DIA based assay (**Figure 4 E**). Inspecting the protein groups showed that the Stellar MS was readily able to target proteins with plasma concentrations ranging from 440 ug/mL (Ceruloplasmin) down to 6.2 fg/mL (Tubulin folding cofactor D) in a single 60 SPD assay design ((**Figure 4 F**) 22 May 2024 www.proteinatlas.org/humanproteome/blood+protein/proteins+detected+in+ms)

### ^15^ N labeled proteins in targeted proteomics

Our data demonstrated that the novel Stellar MS is capable of targeting a large number of peptides in short gradients. To move towards absolutely quantitative assays, we next explored ^15^N labeled proteins standards from our previously discovered liver biomarker panels^19^. Seven of these proteins were recombinantly produced by Absea Biotechnology at sufficient scale for a very large number of assays (**Experimental Procedures**).

This enables the design of targeted assays based on whole ^15^N labeled protein panels where we target multiple peptides at the same time for each protein, significantly increasing the number of targets per assay compared to labeled peptide standards. Unlike C terminal stable isotope labeled synthetic peptides that have a fixed mass shift for every peptide, the mass shift of ^15^N labeled peptides depends on their amino acid (aa) composition and length ranging from d4-d16 in tryptic digests (**Figure 5 A**). The tryptic peptides targeted for the selected proteins cover the range of typically observed mass shifts (**Figure 5 A**, red crosses). Since every aa is labeled, both b and y fragment ions are shifted based on their nitrogen content and charge (**Figure 5 B**).

**Figure 5:**
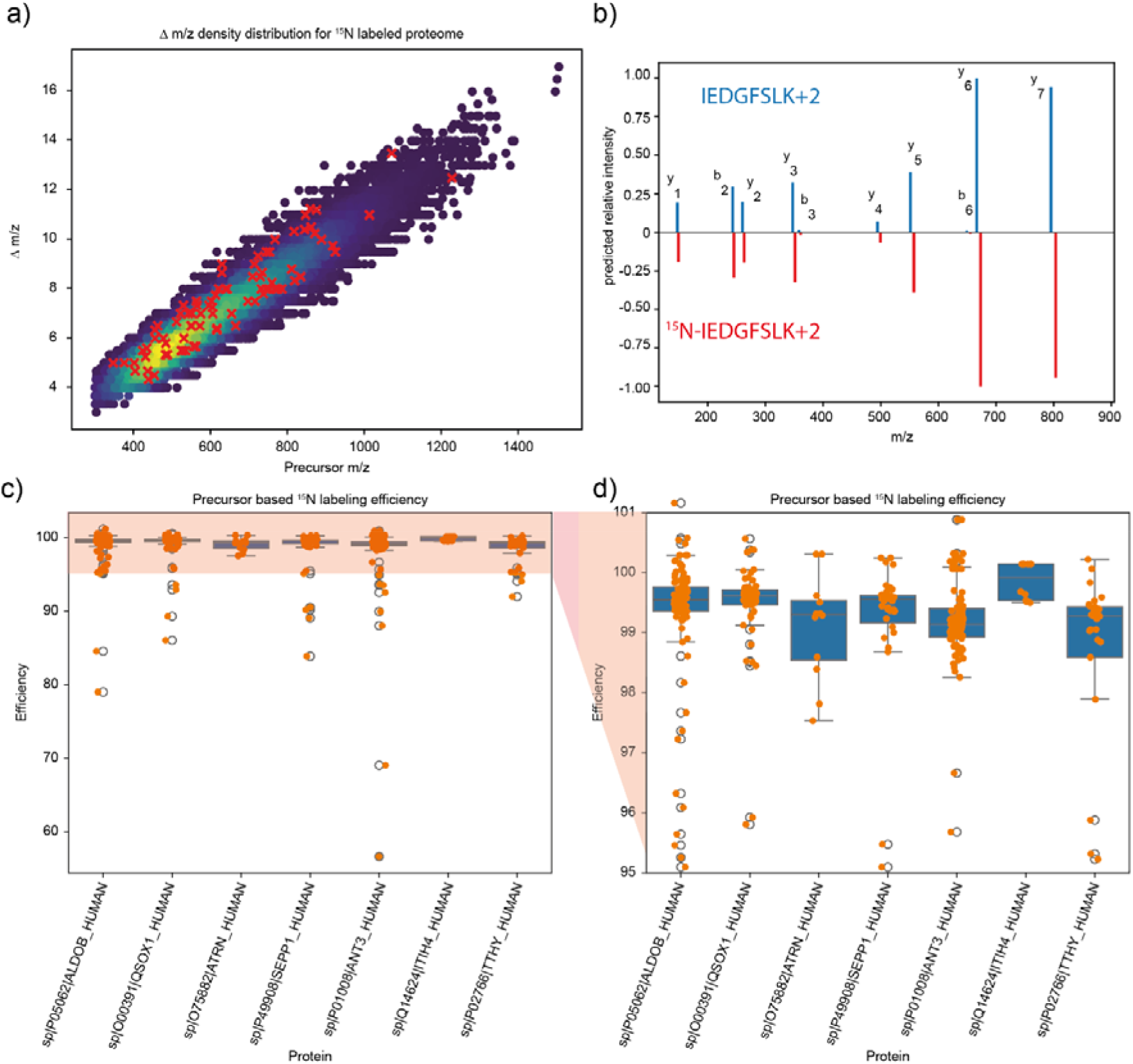
Characterization of ^15^ N Labeled Protein Standards. A Distribution of mass shifts for a ^15^ N labeled total human proteome. The histogram shows the range of mass shifts calculated for tryptic peptides, reflecting the mass shift due to 15N across different amino acid compositions and lengths. B MS2 mirror plot comparing the calculated fragmentation patterns of a heavy ^15^N labeled peptide and its light counterpart. The mirror plot illustrates the individual mass shifts of the between the labeled and unlabeled peptides. C Estimated labeling efficiency of selected proteins based on their precursor isotope envelopes. The graph shows the experimentally measured degree of ^15^N incorporation for each protein, calculated by the ratio of monoisotopic (M0) and M-1 peaks, with a linear regression applied to predict labeling efficiency (Experimental Procedures). D Detailed view of labeling efficiency for specific proteins within the selected panel, confirming over 99% labeling efficiency for all analyzed proteins. This high efficiency ensures accurate quantification and reduces potential interferences in mass spectrometric analysis.

We evaluated the degree of ^15^N labeling for our target panel based on their precursor envelope. Incomplete labeling would have led to highly complex MS2 spectra with distorted fragment ion ratios depending on the position of missed labels but this was not the case. A characteristic feature of isotope contents are the ratios between the monoisotopic mass peak (M0) and its counterparts, here specifically the M-1 peak, in which only one atom was exchanged with a heavier isotope. Such an extracted isotope envelope is illustrated for the peptide IEDGFSLK in Supplementary Figure 1. We calculated the M-1/M0 ratios for every peptide by predicting MS1 envelopes for different simulated labeling efficiencies and thereby accurately determined the labeling efficiency of every precursor using a linear regression as introduced previously^20^ (**Supplementary Figure 1**). Based on this procedure we estimate a degree of labeling of more than 99% for all of our proteins (**Figure 5 C, D**).

### Designing a targeted acquisition scheme for liver diseases using ^15^N labeled protein standards

With a set of high-quality protein standards in hand we designed a targeted acquisition scheme for the measurement of the ^15^N labeled protein panel and their light counterparts. Targeting the standard mix alone at 25 ng each in a 60 SPD gradient, we achieve a median CV of 8.5% across 123 targeted peptides with an average of 11.6 data points per peak (**Figure 6 A, Supplementary Figure 2**). This CV changed only slightly to 8.3% even in a 100 SPD gradient, while the number of data points was reduced to 6 (**Supplementary Figure 2**). The limits of detection (LOD) and limits of quantification (LOQ) were as expected with the majority of LODs and LOQs being below the 0.1 ng dilution level (**Supplementary Figure 3**).

**Figure 6:**
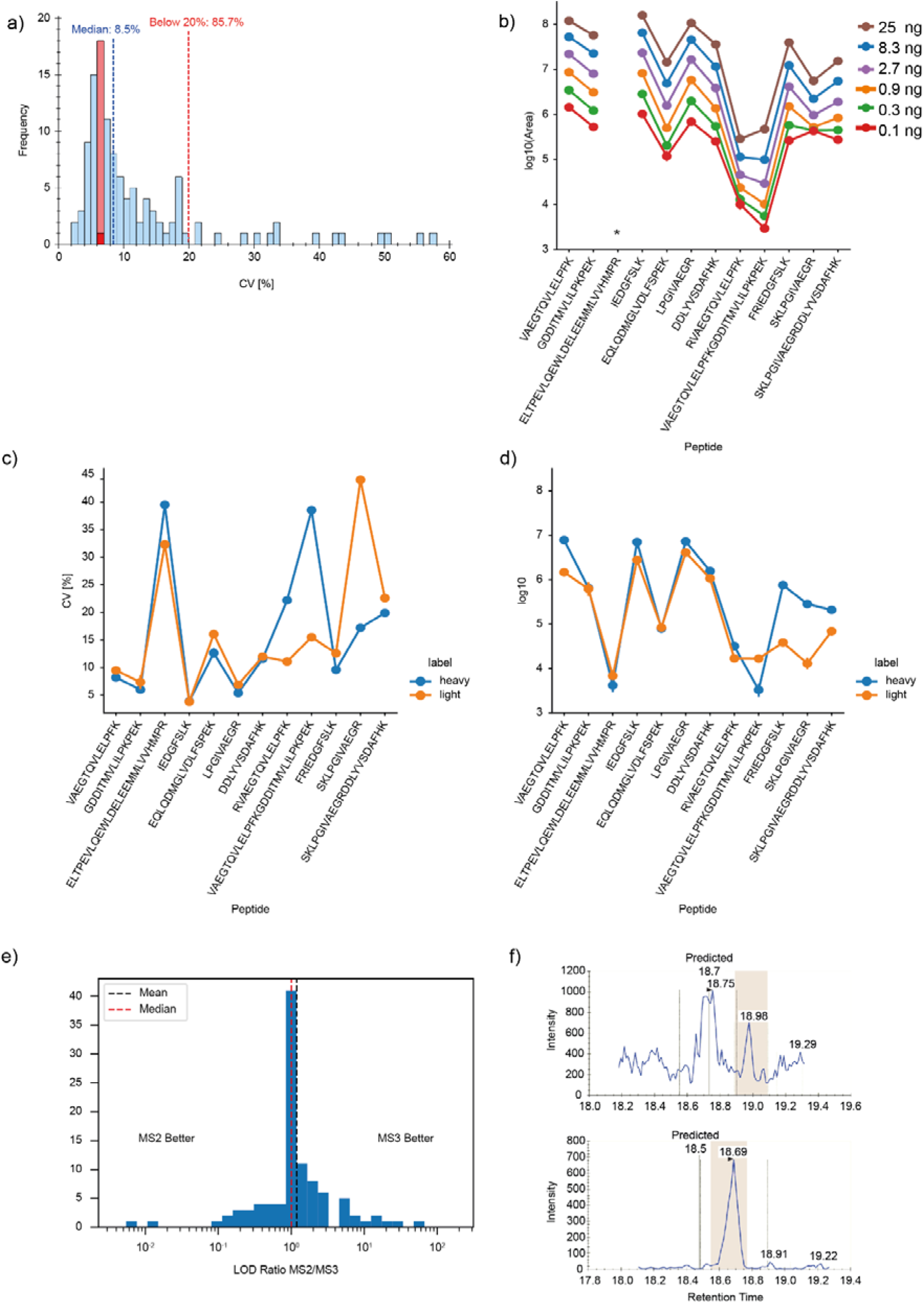
^15^N targeting of liver panel proteins. A Coefficients of variation (CVs) for ^15^N labeled peptides in the liver protein panel measured by itself. The CVs demonstrate excellent reproducibility of measurements across the targeted peptides. B Peptide area-based signals for peptides along the antithrombin III (ANT3) sequence at different concentrations. This graph shows the consistent detection of peptide signals corresponding to varying levels of ANT3. C CVs for light (orange) and heavy (blue) targeted ANT3 peptides in human plasma. CVs are consistent for light and heavy peptides except for those containing missed cleavage sites. D Peptide area-based signals for light (orange) and heavy (blue) targeted ANT3 peptides in human plasma. Intensity traces are consistent except for peptides containing missed cleavage sites. E Improvement in limits of detection (LOD) using MS3 targeting compared to MS2. The plot demonstrates the enhanced sensitivity achievable with MS3 analysis for about half of the peptides. F Comparison of MS2 and MS3 signal intensities for a specific peptide, showing the increase in signal specificity with MS3 analysis.

Upon closer inspection of the peptides of each protein, their different characteristics such as ionizabilities were reflected in the reproducibility of the mean area measured over several replicates leading to a characteristic protein intensity signature along the protein sequence. This signature shifts up and down as expected when varying protein amount (**Figure 6 B**).

These protein intensity signatures were largely recapitulated when we targeted the respective peptides in a plasma proteome sample where we had spiked in the labeled protein mixture in a fixed amount (1 ng/protein). For some of the peptides in this experiment light and heavy peptide pairs deviate from each other both in terms of area as well as CV trends (**Figure 6 C, D**). We attribute this to the fact that we had digested the labeled protein mixture and the plasma proteome separately from each other, mixing them only right before measuring. Interestingly, the affected peptides contained at least one missed cleavage site, implicating differential digestion efficiencies as a contributing factor. In another example, a closer inspection of an outlier peptide in terms of heavy to light ratio revealed that the corresponding protein sequence position was an annotated phosphorylation site (**Supplementary Figure 4**). Such a phosphorylation event would of course not be present in the recombinant (heavy) version of the peptide.

The dual linear ion trap in the Stellar MS allows for efficient MS3 targeting of selected transitions. Applying this to the set of ^15^N labeled peptides did improve the limits of detectability in many cases (**Figure 6 E**), while overall the LODs and LOQs stayed similar. We speculate that this is due to better signal for MS2 (**Figure 6 F**, upper) and better signal to noise in MS3 (**Figure 6 F**, lower) which plays out differently for different peptides. Thus, a mixed assay based on MS2 and MS3 targeting could further improve the assay performance.

## Discussion

In this study, we developed and described a strategy for designing targeted proteomics assays using the novel hybrid high-speed mass spectrometer, Stellar MS, based on discovery DIA data from the Orbitrap Astral. We demonstrated how protein marker panels identified in discovery studies can be translated into targeted assays, enabling parallel targeting of peptides that span entire protein sequences in a native plasma digest and their ^15^N labeled counterparts.

Our results show that a targeted assay can be designed and implemented reliably within a few steps, significantly reducing development time to just a few days. This streamlined process is advantageous not only for clinical assay development but also for the rapid validation of discovery studies. Findings from established and new studies could readily be combined and validated in a targeted manner and this could even be extended to biomarker candidates resulting from non-MS technologies. We have found that biological relevant proteins often display very small changes in abundance between patient groups in clinical cohorts. In our case, only around 2% of the fibrosis-associated plasma proteins changed more than 1.5 fold in early fibrosis ^12^. Stellar MS can be used to verify those results, and with suitable standards, to absolutely quantify proteins of interest. This capability provides new perspectives on plasma proteomics experiments, allowing the cross-validation of clinical results alongside research findings. Stellar MS has the potential to replace numerous single-protein biomarker assays, targeting and quantifying a wide range of protein concentrations, from hundreds of µgrams per mL to picogram per mL in a single run. This range includes many protein markers in routine clinical use. A report submitted in parallel compared the Stellar MS to a triple quadrupole MS and reported up to 10 fold better LOQs^21^.

We also showed that peptide identification lists can be transferred between machine platforms. Thanks to platforms like MassIVE or PRIDE, research laboratories should be able to use the search results and raw files obtained on high resolution platforms to readily develop targeted assays for their own samples. Beyond plasma proteomics, this has great potential for increasing accessibility of technically challenging fields like single cell proteomics to a broader community by using Stellar to target selected proteins or pathways based on data from a different laboratory. However, for this endeavor the accurate prediction of high-quality targetable peptides still has to be improved.

Targeting ^15^N labeled protein panels on Stellar MS showed great promise. This labeling strategy provides whole protein signatures instead of single peptide ratios, increasing comparability between samples. In principle, this could enable the study of proteoforms including PTMs and mutations as the selective presence or absence of specific peptides can be used as a diagnostic feature, identified by inconsistent heavy to light peptide ratios. The whole protein signature can additionally serve as a quality indicator for different sample preparation steps. Targeting more than one peptide per protein at a time also increases the range in which a protein can be accurately quantified^13^. For low abundance proteins, only peptides over a certain intensity range would be used to reduce the influence of signal to noise ratios at the detection limit. Some examples in which this concept could be relevant are the measurement of the C-reactive protein (CRP), which can range from a few milligram per liter up to hundreds of milligrams in patients with inflammatory conditions, or alpha-fetoprotein (AFP) going from low nanograms up to micrograms in patients with hepatocellular carcinoma ^22^.

Overall, we have showcased the capabilities and potential of Stellar MS, a hybrid high-speed mass spectrometer, to advance targeted proteomics. This technology facilitates the rapid and easy transfer of knowledge from cohort-based discovery studies to clinical practice, bringing us closer to bridging the gap between research and patient care.

## ABBREVIATIONS

ACN: acetonitrile
ALD: Alcohol related liver disease
CAA: Chloroacetic acid
dda: data-dependent acquisition
dia: data-independent acquisition
FA: formic acid
GPF: Gas phase fractionation
HPA: Human protein atlas
LIT: Linear ion trap
LOD: Limit of detection
LOQ: Limit of quantification
MeOH: Methanol
MRM: Multiple reaction monitoring
NCE: Normalized collision energy
PRM: Parallel reaction monitoring
SPD: samples per day
SRM: Selected reaction monitoring

## Acknowledgements

We thank our colleagues at the Max Planck Institute of Biochemistry for help and fruitful discussions. Fan Liu and Tao Chen of Absea Biotechnology are acknowledged for insightful discussions and reagents. This study was supported by the Max Planck Society for Advancement of Science, the European Union’s Horizon 2020 research and innovation program under grant agreement No 874839 (ISLET) and by the Bavarian State Ministry of Health and Care through the research project DigiMed Bayern (www.digimed-bayern.de). M.W., V.A and S.S. acknowledge support from the International Max Planck Research School for Life Sciences – IMPRS-LS.

## Data availability

The raw mass spectrometry data have been deposited in the public proteomics repository MassIVE for reviewer access. This data will be made public upon acceptance of the manuscript.

## Supplemental data

This article contains supplemental data.

## Author contributions

M.W., P.M.R., J.B.M.R., V.A., L.N., P.L., S.H., C.C.J., and M.M. conceptualized and designed the study.

M.W., P.M.R., T.H designed and performed experiments. M.W., S.S and P.M.R. analyzed the data.

M.W., P.M.R, and M.M. wrote the original manuscript draft. All authors read, revised and approved the manuscript.

## Supplements

### Supplementary Tables

**Supplementary Table 1:**
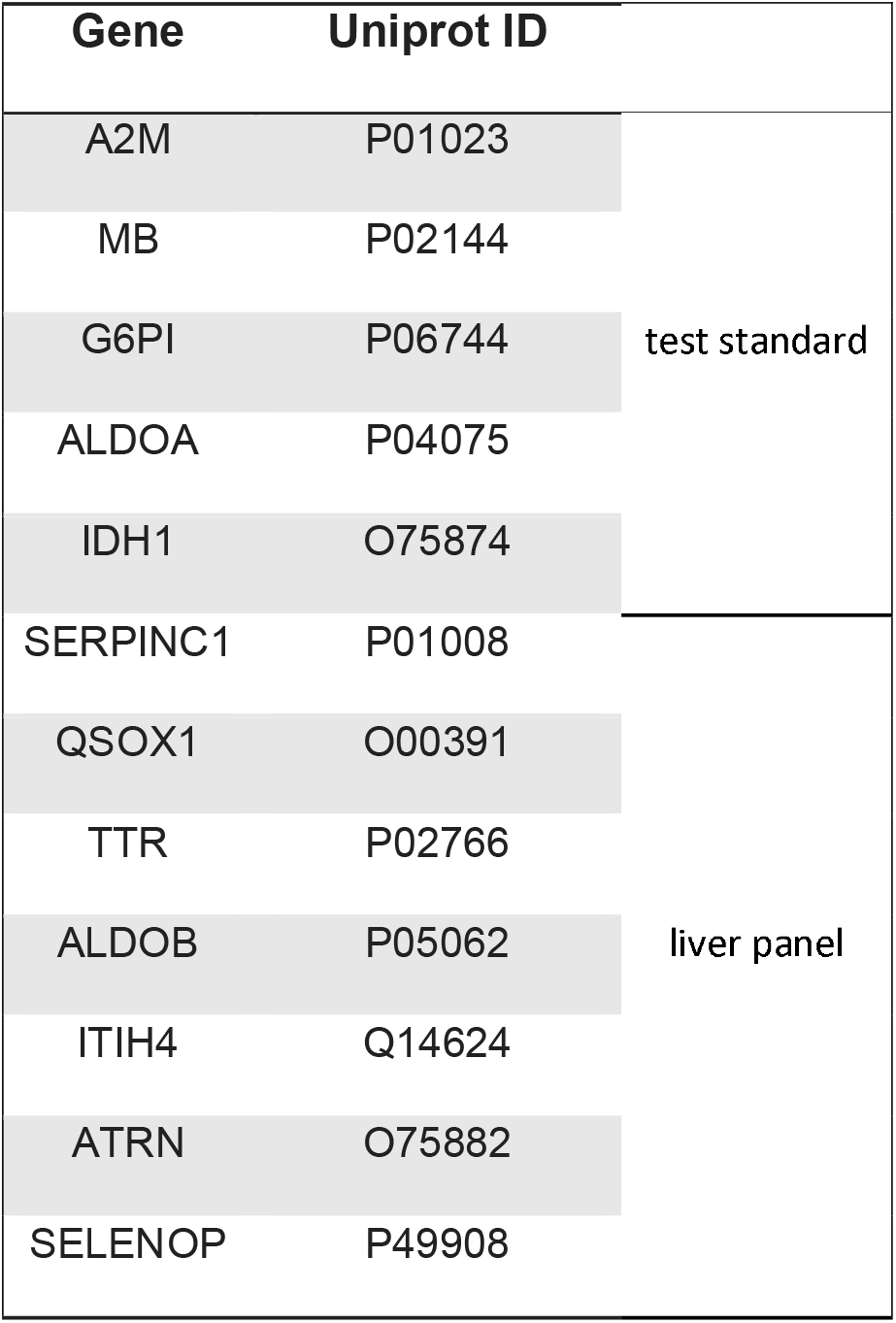
15N labeled protein panel.

| Gene | Uniprot ID |  |
| --- | --- | --- |
| A2M | P01023 | test standard |
| MB | P02144 |  |
| G6PI | P06744 |  |
| ALDOA | P04075 |  |
| IDH1 | O75874 |  |
| SERPINC1 | P01008 | liver panel |
| QSOX1 | O00391 |  |
| TTR | P02766 |  |
| ALDOB | P05062 |  |
| ITIH4 | Q14624 |  |
| ATRN | O75882 |  |
| SELENOP | P49908 |  |

### Supplementary Figures

**Supplementary Figure 1:**
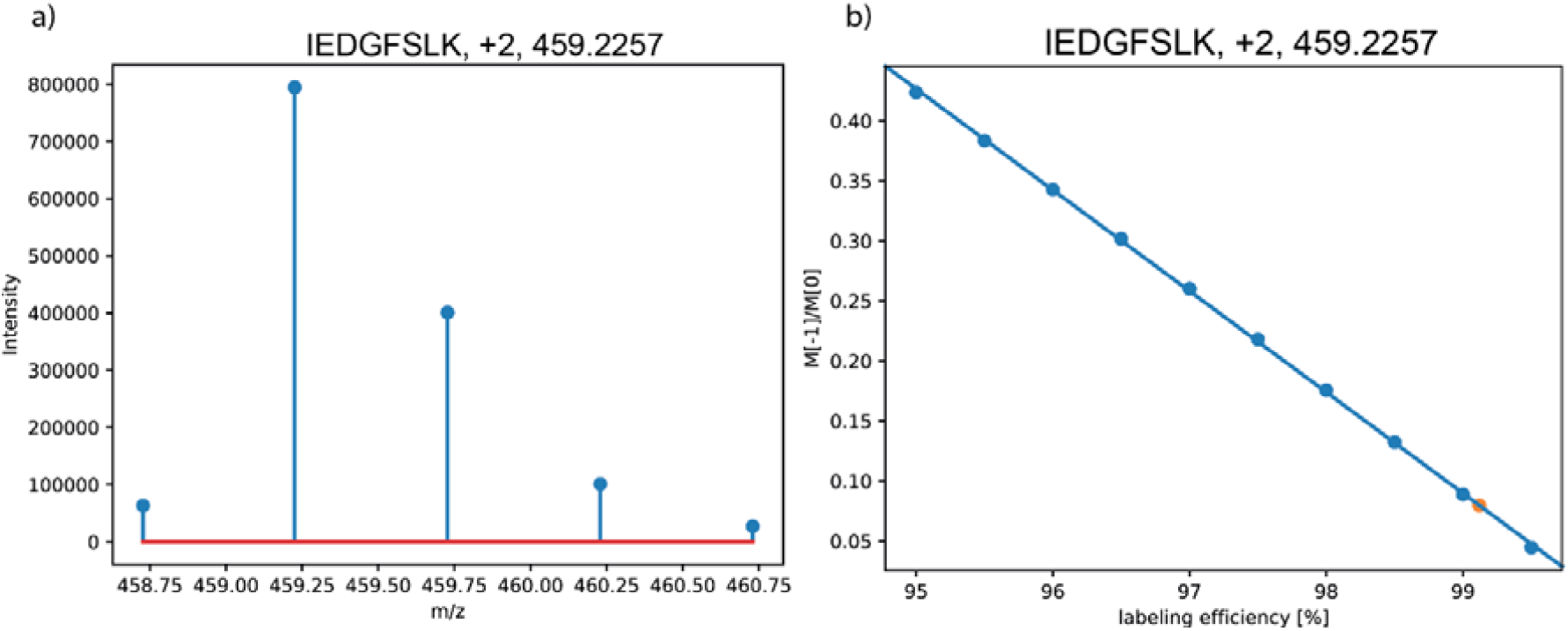
15N labeling efficiency estimation using extracted precursor envelopes. A) Extracted precursor envelope from a DDA acquisition of 15N labeled IEDGFSLK, +2. B) M-1/M0 ratios for different simulated labeling efficiencies (blue) and the extracted M-1/M0 ratio for IEDGFSLK, +2 (orange). The labeling efficiency of IEDGFSLK is estimated using a linear regression. This is repeated for all extracted precursor envelopes to estimate an overall labeling efficiency.

**Supplementary Figure 2:**
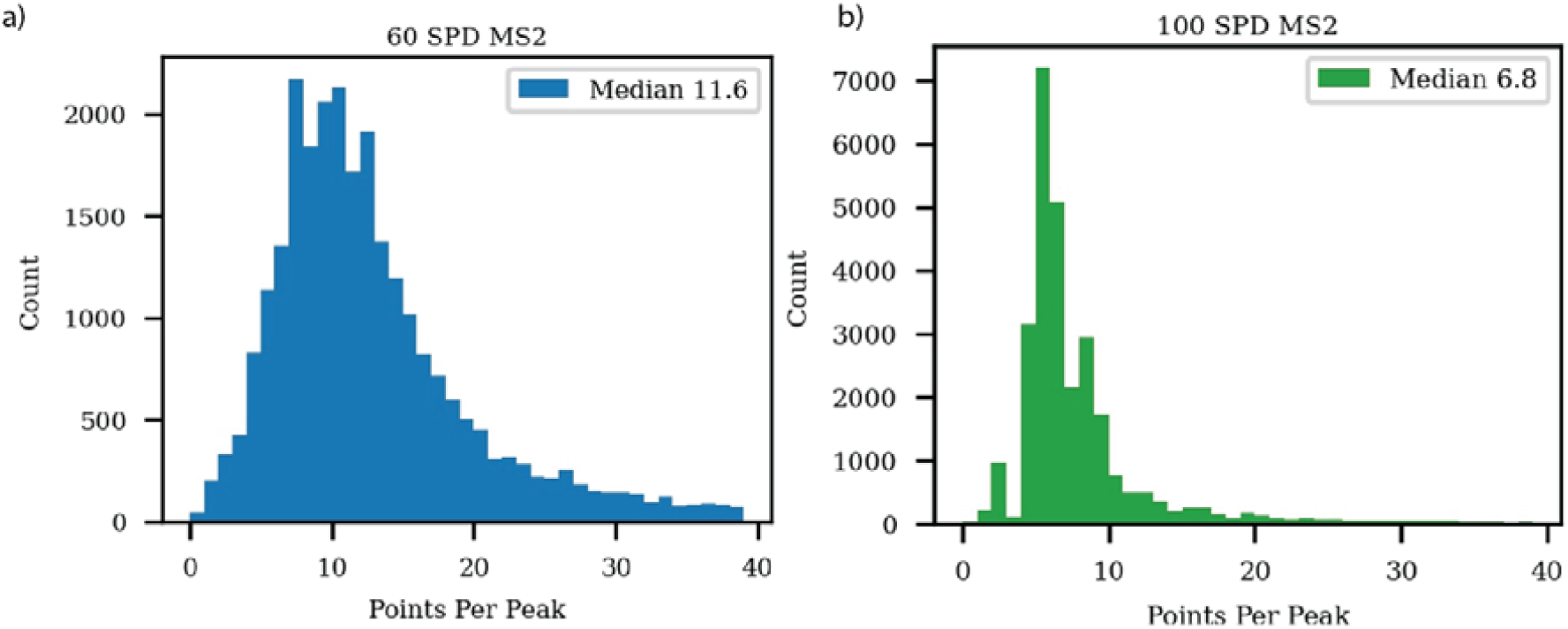
Data points per peak for 15N labeled protein targeting assays. A)Histogram of measured datapoints per peak for 60 SPD MS2 based targeted assay of 15N labeled protein biomarker panel. B)Histogram of measured datapoints per peak for 100 SPD MS2 based targeted assay of 15N labeled protein biomarker panel.

**Supplementary Figure 3:**
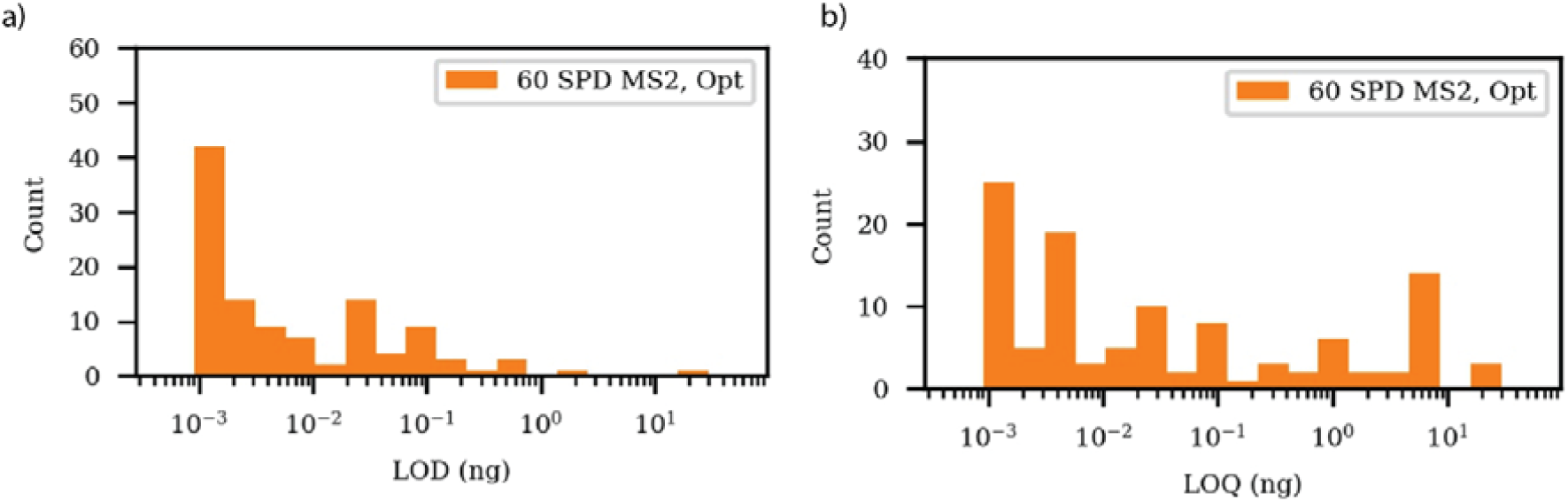
LODs and LOQs of 15N labeled targeting assays. A)Histogram of LODs for each precursor targeted in the 15N labeled targeted biomarker panel. Results are shown for a purely MS2 based assay. B)Histogram of LOQs for each precursor targeted in the 15N labeled targeted biomarker panel. Results are shown for a purely MS2 based assay.

**Supplementary Figure 4:**
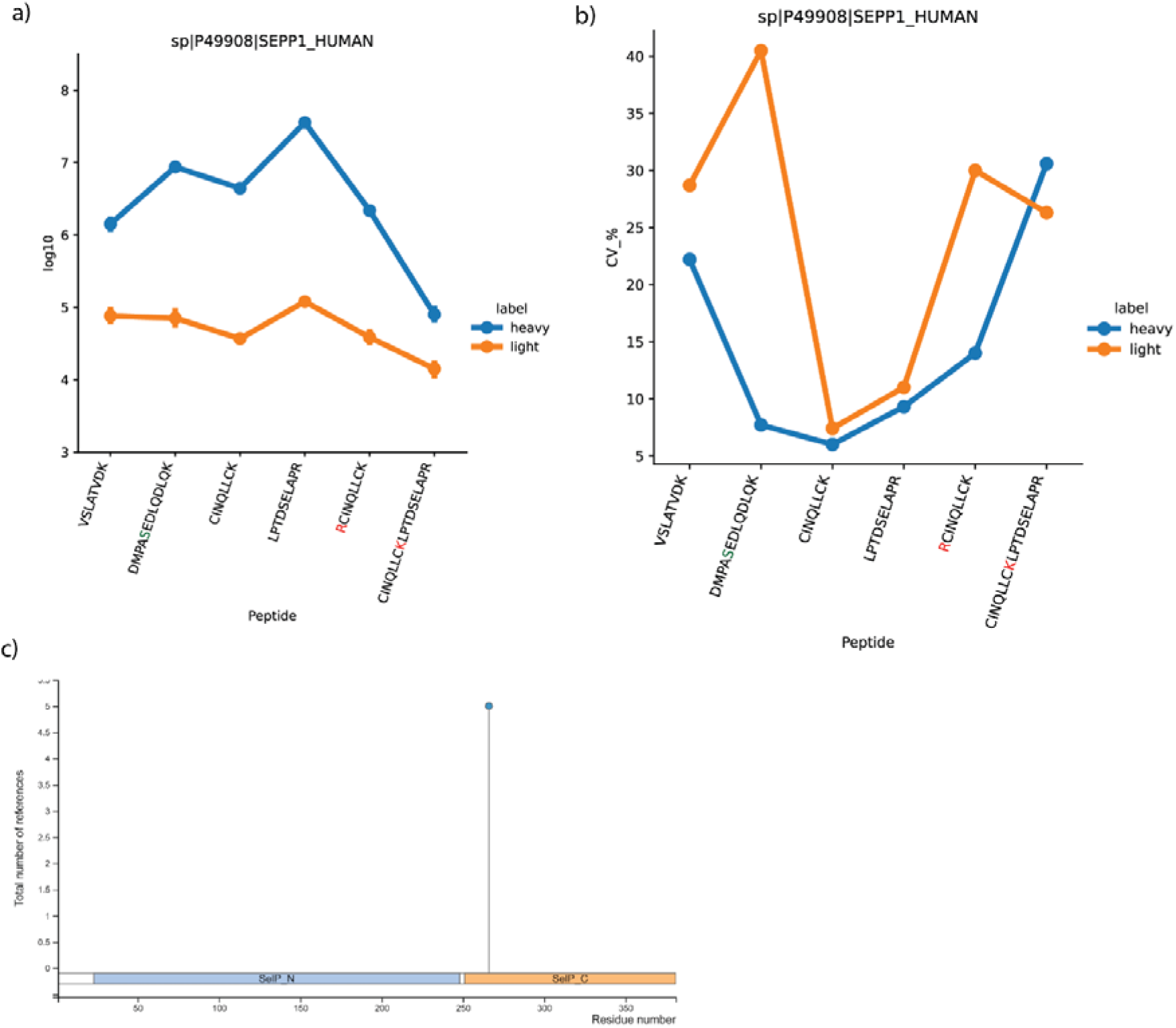
15N labeled targeting of SELENOP. A)Peptide area-based signals for light (orange) and heavy (blue) targeted SELENOP peptides in human plasma. Intensity traces are consistent except for peptides containing missed cleavage sites. B)CVs for light (orange) and heavy (blue) targeted SELENOP peptides in human plasma. CVs are consistent for light and heavy peptides except for those containing missed cleavage sites (red letters) and the peptide potentially carrying a phosphorylation (green letter). C)Post translational modification sites annotated on PhosphoSitePlus. Modifications are filtered for being reported in at least 5 different references with one phosphorylation remaining. (https://www.phosphosite.org/proteinAction?id=22211&showAllSites=true, 24 May 2024)

